# EGFR amplification and PI3K pathway mutations identify a subset of breast cancers that synergistically respond to EGFR and PI3K inhibition

**DOI:** 10.1101/2025.06.03.657674

**Authors:** David J. Wisniewski, Donna Voeller, Yonit A. Addissie, Sachin Kumar Deshmukh, Sharon Wu, Maryam B. Lustberg, Darawalee Wangsa, Danny Wangsa, Kerstin Heselmeyer-Haddad, Yoshimi Endo Greer, George W. Sledge, Stanley Lipkowitz

**Affiliations:** Women’s Malignancies Branch, Center for Cancer Research, National Cancer Institute, Bethesda MD; Caris Life Sciences, Phoenix AZ; Yale Cancer Center, New Haven CT; Genetics Branch, Center for Cancer Research, National Cancer Institute, Bethesda MD

**Keywords:** epidermal growth factor receptor amplification, PI3Kinase pathway mutations, triple negative breast cancer

## Abstract

EGFR family receptor tyrosine kinase signaling is commonly dysregulated in cancer by amplification or activating mutations. Although studies have investigated dual EGFR/PI3K inhibition in breast cancer, they have not determined biomarkers which predict success. We present evidence of a patient subset with EGFR amplification and PI3Kinase pathway mutations in breast cancer which can be synergistically targeted by dual EGFR/PI3K inhibition. This study identified that EGFR amplification occurs in approximately 1-5% of breast cancer patients with shorter overall survival compared to unamplified patients. Up to 71% of EGFR amplified tumors have activating mutations in the PI3K pathway. Dual EGFR/PI3K inhibition more dramatically reduced mTOR and AKT signaling in BT20 and MDA-MB-468 cells which both have EGFR amplification and PI3K pathway activating mutations, compared to control cells. Dual inhibition synergistically reduced cell viability and increased apoptosis in MDA-MB-468 and BT20 compared to control. Single agent therapy in a BT20 xenograft model reduced tumor volume, however only the combination statistically significantly reduced tumor volume compared to control. We conclude that EGFR amplification with co-incident PI3K pathway mutations are driver mutations in a subset of breast cancers and present a subgroup of breast cancers that are more likely to respond to dual targeted therapy.

## Introduction

The ErbB family is comprised of four receptor tyrosine kinases, Epidermal Growth Factor Receptor (EGFR), HER2 (ErbB2), HER3 (ErbB3) and HER4 (ErbB4). Upon ligand binding, the receptors hetero or homodimerize allowing them to activate downstream signaling, including phosphoinositide 3-kinases (PI3Ks) or mitogen-activated protein kinases (MAPKs), resulting in increased proliferation, inhibition of apoptosis, migration, invasion, and other pro-tumor signaling (1–4). Mutation or overexpression of these family members are common drivers of various cancer types (5). For example, HER2 amplification is a driver in 12-20% of breast cancers, correlates with poor survival, and has been effectively targeted to improve outcomes in patients with HER2 amplified breast cancer (6–9).

EGFR can be aberrantly activated through mutations or amplification/overexpression, which drives pro-tumor effects in cancers such as non-small cell lung cancer, pancreatic cancer and head and neck cancer, leading to approval of EGFR targeted therapies including tyrosine kinase inhibitors (TKI’s) gefitinib, erlotinib, afatinib, dacomitinib, osimertinib, and vandetanib, and the monoclonal antibodies cetuximab, panitumumab and necitumumab (1, 2, 10–13). EGFR is often expressed in basal triple negative breast cancer (TNBC), and is associated with tumor size, poor differentiation, worse relapse-free and overall survival, and shorter metastasis free survival (14, 15). Depending on the study, EGFR gene amplification has been observed in 0.8- 14% of breast cancer patients (16–19). Mutation of EGFR in breast cancer can range from approximately 2-11% (20). Relatively small studies looking specifically at metaplastic breast carcinomas have observed amplification of EGFR in 33-37% of patients (21, 22). Despite these data, clinical trials investigating EGFR inhibitors have not shown great promise in breast cancer, with studies finding partial response rates of 0-6% (16, 20, 23–25). These studies were typically in heavily pretreated patients, and patients were not stratified by EGFR amplification status.

Although previous studies in various *in vitro* cancer models have evaluated EGFR and PI3K inhibitor combination, these studies did not investigate the role of EGFR amplification in addition to PI3K mutation status as biomarkers for drug synergy (26–37). Further, they did not characterize a potential patient population which would most benefit from this treatment regimen. Here, we evaluated the frequency and subtype distribution of EGFR amplification, the clinical outcomes of patients with EGFR amplified tumors, the frequency of coincident activating mutations in the PI3K pathway and studied the effects of dual inhibition of EGFR and the PI3K pathways in TNBC cell line models in vitro and in vivo. Our data provides evidence that breast cancers with EGFR amplification and PI3K pathway activating mutations may be more likely to respond to therapies targeting these pathways.

## Methods

### Materials

Z-VAD-FMK (S7023), Erlotinib (S7786), BKM120 (S2247), Afatinib (S1011), Alpelisib (S2814) and sodium carboxymethyl cellulose (S6703) were purchased from Selleck Chemicals. Fetal bovine serum (FBS) (631106) was obtained from Takara Bio. 4-20% precast polyacrylamide gels (5671094) were purchased from Bio-Rad. Immobilon PVDF transfer membrane (IPVH00010), mini protease inhibitor cocktail (11836153001) and propidium iodide (P4864) were purchased from Millipore-Sigma. PBS (21-031-CV), EGF (354042) and Phenol Red-Free LDEV-Free Matrigel (356237) was purchased from Corning. Sodium orthovanadate (P0758S) was from Fisher Chemicals. NP40 Cell Lysis Buffer (FNN0021), Penicillin- Streptomycin (15140–122), tissue extraction reagent (FNN0071) and RPMI 1640 medium (11875–093) was obtained from ThermoFisher Scientific.

### Antibodies

Anti-phospho-HER3 Y1289 (4791S), anti-HER3 (12708S), anti-EGFR (2232), anti-PTEN (9559S), anti-phospho-EGFR Y1068 (2234), anti-phospho-AKT S473 (9271S), anti-AKT (9272S), anti-phospho-MAPK T202/Y204 (9101), anti-phospho-P70S6K T389 (9234S), anti- P70S6K (9202S), anti-phospho-S6 S235/236 (4856S), anti-S6 (2217S) and anti-PI3K (4249S) were purchased from Cell Signaling Technology. anti-HSC70 (sc-7298) and anti-ERK2 (D-2, sc-1647) antibodies were purchased from Santa Cruz Biotechnology. Goat Anti-Mouse IgG HRP conjugate (172-1011) and Goat Anti-Rabbit IgG HRP conjugate (1721019) were purchased from BioRad.

### Cell Lines

MDA-MB-468, BT20, MDA-MB-231 were obtained from ATCC, maintained in RPMI 1640 medium with 10% FBS and 1% penicillin/streptomycin, and grown at 37°C, 5% CO_2_. These cell lines were recently authenticated and tested for mycoplasma contamination.

### Caris data

#### Next-generation sequencing (NGS)

NGS was performed on genomic DNA using the NextSeq (592-whole gene targets; Agilent Technologies, Santa Clara, CA, USA) or NovaSeq 6000 (>700 genes at high coverage/read- depth and >20 000 genes at lower depth; Agilent Technologies, Santa Clara, CA, USA) platforms (Illumina, Inc., San Diego, CA, USA) as described previously (38). The copy number alteration (CNA) of each exon is determined by normalizing the sequencing depth of each exon divided by the average sequencing depth of the sample and comparing it to the pre-calibrated mean of normalized values in the training data (The mean values are re-calibrated every 60 days with up to 10,000 samples). A gene was defined as amplified by having segment level of copies ≥4.5.

#### Survival analysis

Caris CodeAI™ clinico-genomic database containing insurance claims data were used to calculate real-world overall survival (OS) from tissue collection to last contact. Kaplan-Meier curves were generated to calculate molecularly defined patient cohorts as previously described (39, 40).

#### cBioPortal data

Datasets from TCGA (Cell 2015), METABRIC (Nature 2012 and Nature Communications 2016) and MSKCC (Cell 2018) were searched for EGFR amplification, as well as mutations in PIK3CA, PIK3R1, PIK3R2, PIK3R3, PTEN, AKT1, AKT2, AKT3, RPS6KB1, RPS6KB2 and RPS6 in cBioPortal (cbioportal.org) (41–43). Amplification was defined by GISTIC2.0 score = 2 (CN ≥4-5) (44).

### Western Blot

Cell lysates were collected, prepared and immunoblotted as previously described, and repeated at least 3 times (45). Densitometric analysis of band intensities was calculated using ImageJ and data are presented as average ± SEM.

### Fluorescence *in situ* Hybridization (FISH)

To prepare metaphase chromosomes, cells were treated with 0.02 mg/ml Colcemid (Invitrogen, Grand Island, NY) for one hour, suspended in a hypotonic solution for 20 minutes, fixed using a methanol/acetic acid mixture (1:3 ratio) and dropped onto slides in a cytogenetic drying system (Thermotron, Holland, MI). Dual color FISH was performed with custom probes for EGFR and centromere 7 from Cytotest (Rockville, MD), labeled with Dyomics dyes (Jena, Germany).

Slides were pepsin treated, washed in 1x PBS, dehydrated in an ethanol series, and left to air-dry. Co-denaturation was performed at 72°C for 2 minutes, followed by overnight hybridization at 37°C. The slides were detected in 2× SSC/0.3% Nonidet P-40 for 2 minutes at 48°C, and 2× SSC/0.1% Nonidet P-40 for 1 minute at room temperature and were mounted with DAPI antifade solution (Vector Laboratories, Burlingame, CA). Images were captured using a Leica DM-RXA fluorescence microscope (Leica, Wetzlar, Germany) equipped with a 40 X objective and custom optical filters (Chroma, Bellows Falls, VT).

### Sanger Sequencing

Sanger DNA sequencing of genomic DNA from the MDA-MB-231, MDA-MB-468 and BT20 cell lines for PIK3CA mutations was performed using the Big Dye Terminator v.1.1 Cycle Sequencing Kit (Applied Biosystems, CA) according to the manufacturer’s specifications and run on an ABI 3130xl or 3730 Genetic Analyzer (Applied Biosystems, CA) at the CCR Genomics Core at the National Cancer Institute.

### Cell Death

Cell death was determined by CytoTox-Glo (G9291) as directed by Promega or by propidium iodide uptake. Propidium iodide (PI) staining, imaging and quantification was performed using the BioTek Cytation 1 Imaging Reader from Agilent (Santa Clara, CA) and Gen5 Image Prime 3.14 software (Agilent). For CytoTox Glo, cells were pretreated with 20 µM ZVAD for 1 hour, followed by co-treatment with DMSO, 10 µM erlotinib, 1 µM BKM120 or the combination for 48 hours. Dead cell percentage was then calculated according to the manufacturer instructions. For PI, 5000 cells were plated in a 96-well plate and treated the next day with experimental conditions in 1% FBS RPMI, along with PI (1:3000 dilution). Dead cell percentage was calculated by dividing dead cell number (PI stained) by brightfield cell count number and multiplying by 100 and normalized relative to the Day 0 dead cell percentage. In experiments using ZVAD-FMK, cells were pretreated +/- 20 µM ZVAD-FMK for 1 hour, followed by co- treatment with experimental conditions.

### Cell Viability

10 000 cells were plated in a 96-well white walled plate (3610, Corning) and treated the following day with experimental conditions for 48 hours in 1% FBS RPMI. Viability was determined by CellTiter-Glo 2.0 (Catalog #G9242) as directed by Promega.

### Cell cycle

Cells were plated in 6-well plates (3x10^5^ cells/well) and treated with EGFR and/or PI3K inhibitors for 48 hours. DNA content was measured by flow cytometry as previously described, using a BD LSRFortessa™ (BD Biosciences) flow cytometer and analyzed using FlowJo® software (FlowJo LLC) (46, 47).

### Animal Studies

Sixty female athymic nude mice between 5-6 weeks in age obtained from Charles River Laboratories were housed and treated according to approved NCI-ACUC guidelines and in accordance with an approved NCI Animal Care and Use Committee (IACUC number WMB- 004). 5 million BT20 cells suspended in matrigel (1:1 with PBS) were injected subcutaneously into the mammary fat pad with a 25G needle (Terumo Medical Care Solutions; SS-01T2516). When tumors were 100 mm^3^ the mice were randomized into four groups of 15. The mice were randomized so each group included similar average tumor size, with similar standard deviation. 0.5% sodium carboxymethyl cellulose (CMC) in water (w/v) was used as control, and 20 mg/kg afatinib or 20 mg/kg alpelisib were suspended in 0.5% CMC. Control or 20 mg/kg alpelisib, 20 mg/kg afatinib, or the combination of both was given by oral gavage (1 inch, 22G needle) once a day for five days each week for the duration of the study. Animals were not included in treatment or analysis if the tumors did not reach 100 mm^3^ at the time of randomization. After the first four days of treatment, four mice per group were sacrificed, tumors were excised and proteins were harvested in Tissue Extraction Reagent I (Invitrogen-FNN0071) according to the manufacturers protocol. Tumor caliper measurements and body weight were measured twice a week for the duration of the study, as required by humane endpoints for all mice. Investigators were blinded to the tumor measurements. Tumor volumes were calculated according to the formula V= ½ (length×width^2^). When maximum tumor size for each animal was reached (20 mm in any direction) the mice were humanely euthanized.

### Statistical Analysis

Student’s *t*-test with 2 tailed comparisons assuming equal variance, and one-way or two-way ANOVA were performed as indicated. P-values of ≤0.05 were considered significant. Synergy scores were calculated using SynergyFinder.org (48). *In vitro* experiments consisted of triplicates per treatment group, with at least three experimental repeats performed (with specific repeat numbers indicated in figure legends). *In vivo* experiments contained n=15 per treatment group.

## Results

### Incidence of EGFR amplification in breast cancer

EGFR amplification was observed in 1.2-2.4% of cases in the TCGA, METABRIC and MSKCC databases in cBioPortal (41–43), depending on the dataset (Table 1). The TCGA and METABRIC databases contain mostly early-stage primary breast cancers while the MSKCC dataset is approximately 50% metastatic/recurrent breast cancers. EGFR amplification is more common in invasive ductal carcinomas (IDC) (1.4-2.9%) compared to invasive lobular carcinomas (ILC) (0-0.6%). Further, the highest rate of amplification was in TNBC (3.2-6.7%), ER- (2.8-5.4%) and ER-/HER2+ tumors (1.3-6.5%), whereas EGFR amplification was relatively less common in ER+ breast cancers (0.8-1.4%). Using PAM50 classification, EGFR amplification is the highest in basal (3.7-8.1%) and HER2 enriched (3.9-5.4%) molecular subtypes, compared to luminal A or luminal B subtypes (0.7% and 1.5-1.6%, respectively) (49). We combined the TCGA, METABRIC and MSKCC datasets due the small number of EGFR amplified cases in each and found that there is a statistically significant lower OS in patients with amplified EGFR, with a difference of 53.5 months (111.4 vs 164.6 months) (Fig. 1A).

**Figure 1.**
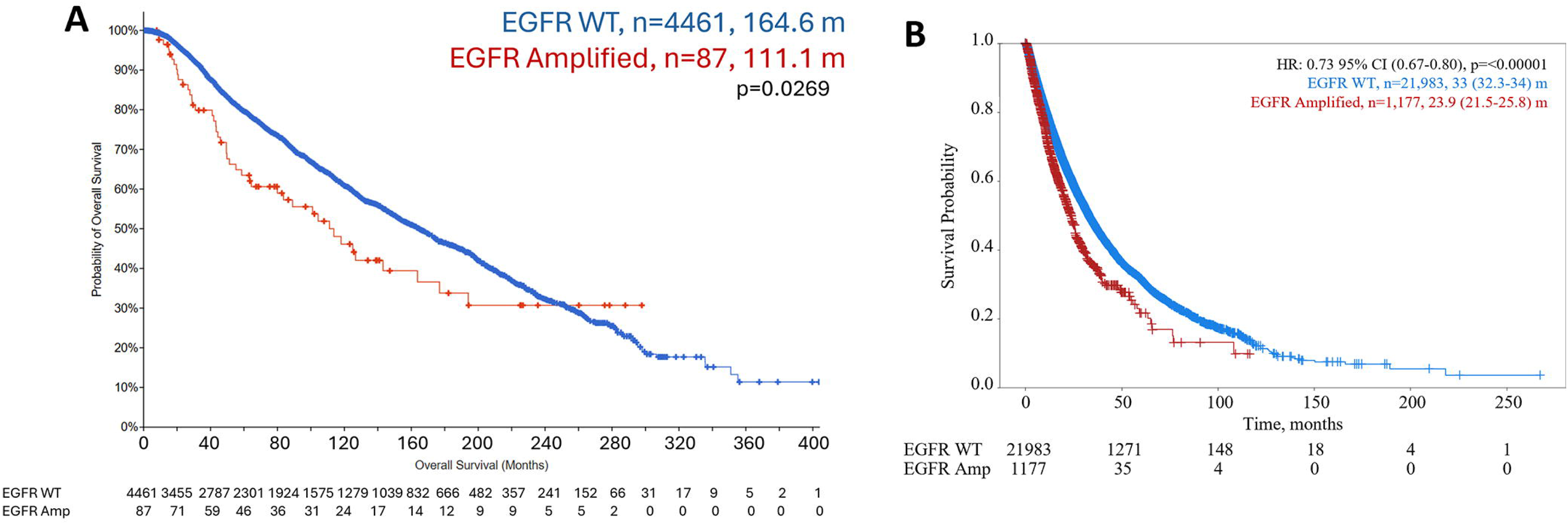
Survival of patients with EGFR amplification in breast cancer. A) Combination of TCGA, METABRIC, MSKCC datasets to analyze overall survival (OS) of patients comparing EGFR amplification (red line) to patients without EGFR amplification (blue line). B) OS of patients from Caris dataset with EGFR amplification (red line) compared to patients without EGFR amplification (blue line).

**Table 1.** Incidence of EGFR amplification in breast cancer. Breast cancer subtype analysis of patients in TCGA, METABRIC, MSKCC and Caris dataset with EGFR amplification. IDC, invasive ductal carcinoma; ILC, invasive lobular carcinoma; nd, not done.

|  | TCGA |  | METABRIC |  | MSKCC |  | Caris |  |
| --- | --- | --- | --- | --- | --- | --- | --- | --- |
| Breast Cancer Subtype | Samples # | EGFR Amplification (%) | Samples # | EGFR Amplification (%) | Samples # | EGFR Amplification (%) | Samples # | EGFR Amplification (%) |
| All Patients | 816 | 14 (1.7%) | 2173 | 52 (2.4%) | 1918 | 23 (1.2%) | 23160 | 1177 (5.1%) |
| Histology |  |  |  |  |  |  |  |  |
| IDC | 489 | 10 (2.0%) | 1660 | 48 (2.9%) | 1473 | 21 (1.4%) | 8411 | 434 (5.2%) |
| Mixed IDC/ILC | 88 | 0 (0%) | 218 | 2 (0.9%) | 89 | 0 (0%) | nd | nd |
| ILC | 127 | 0 (0%) | 172 | 1 (0.6%) | 299 | 0 (0%) | 1311 | 27 (2.1%) |
| Other Histologies | 112 | 4 (3.6%) | 123 | 1 (0.8%) | 57 | 2 (3.5%) | 13438 | 716 (5.3%) |
| ER and HER2 Status |  |  |  |  |  |  |  |  |
| ER+ | 600 | 5 (0.8%) | 1617 | 23 (1.4%) | 1512 | 13 (0.9%) | 10802 | 363 (3.4%) |
| ER+/HER2- | 326 | 3 (0.9%) | 1398 | 19 (1.4%) | 1357 | 11 (0.8%) | 9887 | 342 (3.5%) |
| ER Negative | 175 | 8 (4.6%) | 523 | 28 (5.4%) | 327 | 9 (2.8%) | 6312 | 512 (8.1%) |
| HER2 Positive | 121 | 3 (2.5%) | 247 | 9 (3.6%) | 240 | 3 (1.3%) | 1625 | 73 (4.5%) |
| ER+/HER2+ | 89 | 1 (1.1%) | 108 | 1 (0.9%) | 76 | 1 (1.3%) | 916 | 21 (2.3%) |
| ER-/HER2+ | 31 | 2 (6.5%) | 139 | 8 (5.8%) | 79 | 1 (1.3%) | 709 | 52 (7.3%) |
| TNBC | 90 | 6 (6.7%) | 335 | 19 (5.7%) | 251 | 8 (3.2%) | 5335 | 446 (8.4%) |
| Molecular Subtype |  |  |  |  |  |  |  |  |
| Total Profiled | 587 | 10 (1.7%) | 1980 | 47 (2.4%) | nd | nd | 12872 | 611 (4.6%) |
| Luminal A | 307 | 2 (0.7%) | 700 | 5 (0.7%) | nd | nd | 1764 | 51 (2.9%) |
| Luminal B | 122 | 2 (1.6%) | 475 | 7 (1.5%) | nd | nd | 5402 | 153 (2.8%) |
| HER2 Enriched | 51 | 2 (3.9%) | 224 | 12 (5.4%) | nd | nd | 2233 | 173 (7.7%) |
| Basal | 107 | 4 (3.7%) | 209 | 17 (8.1) | nd | nd | 3367 | 230 (6.8%) |
| Claudin Low | nd | nd | 218 | 3 (1.4%) | nd | nd | nd | nd |
| Normal | nd | nd | 148 | 3 (2.0%) | nd | nd | 106 | 4 (3.8%) |

To confirm these data in a larger dataset with clinical annotation, we evaluated the incidence of EGFR amplification (≥4.5 CN) in patients with breast cancer profiled by Caris Life Sciences (Caris), where the majority of patients profiled had metastatic breast cancer (Table 1). EGFR amplification was found in 5.1% of the breast cancers profiled. As in the other datasets, EGFR amplification was more prevalent in IDC (5.2%) compared to ILC (2.1%). EGFR amplification was more prevalent in TNBC (8.4%), and in the molecular HER2 enriched (7.7 %), basal (6.8%) and normal (3.8%) subtypes. The EGFR amplification seen in TNBC in this dataset (8.4%) is similar to previously published data, which found 8.9% EGFR amplification in TNBC (50). OS was statistically significantly shorter by 9.1 months in patients with EGFR amplification compared to those with unamplified EGFR (23.9 vs 33 months; p<0.00001: Fig. 1B). The shorter survival in the Caris dataset is likely due to the patients being metastatic, while the majority of the patients in the dataset for Figure 1A were early-stage patients. This data shows that there is a subset of breast cancer patients with EGFR amplification that is most frequent in TNBC and ER-/HER2 amplified tumors and is associated with a poor prognosis.

Since EGFR amplification is enriched in patients with TNBC or Basal cancers, we investigated the differences in overall survival in patients with EGFR amplification depending on subtype using the Caris dataset (Fig. S1). There was a borderline worse overall survival in patients with EGFR amplification (difference of 1.7 months; p=0.09) in TNBC patients (Fig. S1A). Patients with molecularly defined Basal tumors with EGFR amplification also exhibited a borderline worse overall survival (difference of 1.3 months; p=0.06) (Fig. S1B). There was no significant difference in the survival of HER2 enriched patients with EGFR amplification (Fig. S1C). In contrast, overall survival was statistically significantly worse in patients with ER+/HER2- tumors that had EGFR amplification compared to those with WT EGFR (difference 9.6 months; p<0.001) (Fig. S1D).

### *In vitro* modeling of EGFR amplification and PI3K alterations

To evaluate the role of EGFR amplification in cell models, we used two cell lines with amplified EGFR (MDA-MB-468 and BT20) and one cell line without amplified EGFR (MDA-MB-231). MDA-MB-468 and BT20 have elevated EGFR protein levels as compared to MDA-MB-231 and FISH confirmed the amplification of EGFR in both MDA-MB-468 and BT20, and the lack of EGFR amplification in MDA-MB-231 (Fig. 2A and B). MDA-MB-468 has loss of PTEN protein, which supports previously published studies showing loss of PTEN expression in these cells (Fig. 2A) (51, 52). As previously reported, the BT20 TNBC cell line has two activating mutations in PIKCA (P539R and H1047R), which previously have been shown to be in cis (Fig. 2C) (52–54). MDA-MB-468 and MDA-MB-231 have wildtype PIK3CA. Therefore, MDA-MB-468 and BT20 were the two cell lines with EGFR amplification and PI3Kinase pathway activating mutations, and MDA-MB-231 was the control cell line without amplified EGFR or PI3Kinase pathway mutations.

**Figure 2.**
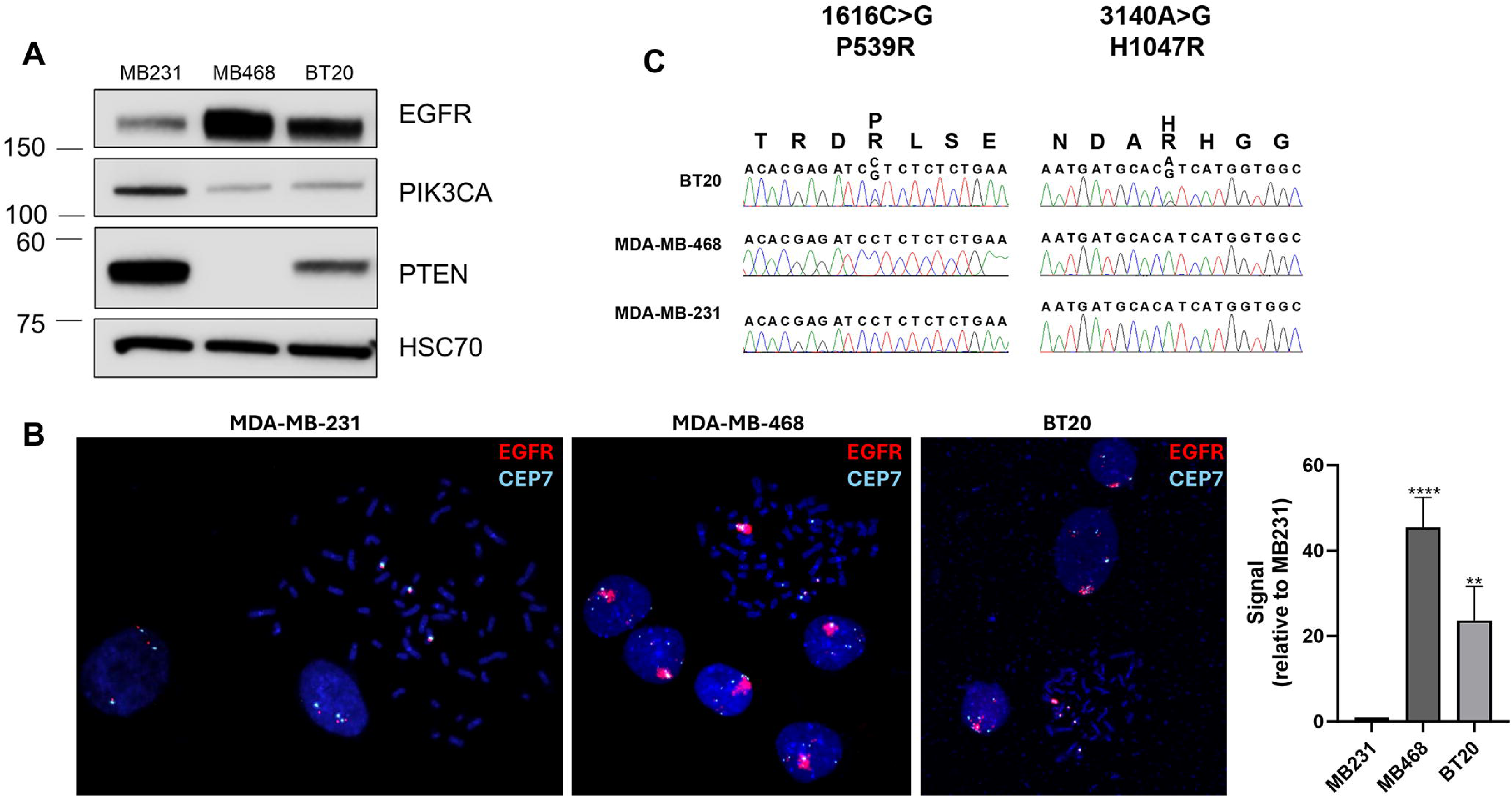
EGFR protein expression and gene amplification, and PI3K mutations in MDA-MB-231, MDA-MB-468 and BT20 cells. A) Western blot showing protein expression of EGFR, PIK3CA, pTEN and HSC70 (loading control) in MDA-MB-231, MDA-MB-468 and BT20 cells. B) FISH analysis showing EGFR (red) and CEP7 (aqua, control) in MDA-MB-231, MDA-MB-468 and BT20 cell lines. Average EGFR signal per nuclei was quantified using ImageJ, and statistical analysis was performed by Student’s t-test, with **** indicating p<0.0001 and ** indicating p<0.01 compared to MB231. C) DNA sequencing for PIK3CA mutation status in BT20, MDA-MB-468 and MDA-MB-231 cell lines.

### PI3K pathway mutations in EGFR amplified breast cancer

Considering the co-incidence of mutations in PI3K signaling in these EGFR amplified cell lines, using cBioPortal we investigated the incidence in patients. In 75 patients with EGFR amplification from the TCGA, METABRIC and MSKCC cohorts, 53 patients (approximately 71%) with EGFR amplified tumors have at least one genetic alteration which can potentially activate PI3K signaling (Table 2 and Fig S2). 36% have an activating mutation in PIK3CA, 5.3% have a loss of PTEN, 12% have mutations in PIK3R1, R2 or R3, and 29.3% have amplification of AKT1, 2, or 3. There are 16% of patients who have amplifications downstream of mTOR (either in ribosomal protein S6 kinase B1 or B2, or ribosomal protein S6). Of note, multiple tumors had two or more alterations in this pathway (Fig. S2).

**Table 2.** PI3K pathway mutations in EGFR amplified breast cancer. Datasets from TCGA, METABRIC and MSKCC combined showing alterations in PI3K pathway genes in 75 patients with EGFR amplification. Dataset from Caris showing alterations in PI3K pathway genes in 1,177 breast cancers with EGFR amplification.

|  | TCGA, METABRIC, MSKCC combined | Caris |
| --- | --- | --- |
| Gene | Potential PI3KPathway Alterations (%) | Potential PI3KPathway Alterations (%) |
| PIK3CA | 36 | 36.5 |
| PIK3R1 | 4 | 4.32 |
| PIK3R2 | 4 | 0 |
| PIK3R3 | 4 | Not Available |
| PTEN | 5.3 | 10.5 |
| AKT1 | 5.3 | 5.62 |
| AKT2 | 6.7 | 0 |
| AKT3 | 17.3 | 0 |
| RPS6KB1 | 8 | Not Available |
| RPS6KB2 | 4 | Not Available |
| RPS6 | 4 | Not Available |
| Patients with at least one PI3K activation | 70.7 | 50.8 |

In the Caris dataset, 50.8% of EGFR amplified tumors have mutation or amplification in the PI3K pathway (including PIK3CA, pTEN, PIK3R1, PIK3R2, or AKT 1, 2 or 3) (Table 2). The largest percentage of alterations are in PIK3CA (36.5%). The Caris dataset does not include alterations in the genes downstream of mTOR, contributing to the difference comparing the sums of Caris and the cBioPortal data. This data confirms that many tumors with EGFR amplification are likely to have co-incident aberrant PI3K signaling.

### EGFR and PI3K dual inhibition significantly reduce downstream signaling in EGFR amplified and PI3K altered breast cancer

Epidermal growth factor (EGF) and tyrosine kinase inhibitor (TKI) treatment showed that EGFR phosphorylation was decreased upon afatinib or erlotinib treatment in both MDA-MB-468 and BT20 cell lines, indicating that the TKI inhibited EGFR signaling upon EGF stimulation (Fig. 3A, Fig. S3A). To determine the effects of EGFR and PIK3CA dual inhibition on steady state signaling, we looked at MAPK, AKT, P70S6K and S6K activation. In the EGFR amplified cell lines MDA-MB-468 and BT20, afatinib significantly inhibited basal MAPK activation, but had no significant effect on MAPK signaling in MDA-MB-231 cells (Fig. 3B and C). Therefore, EGFR amplified cells have activated MAPK signaling downstream of EGFR. Afatanib alone also down regulated AKT signaling in the MDA-MB-468 cells but not in the BT20 or the MDA-MB-231 cell lines (Fig. 3B and D, Fig. S3B). The PIK3CA inhibitor alpelisib significantly down regulated AKT activation in all three cell lines but did not affect pMAPK (Fig. 3B and D, Fig. S3B). Combination of afatanib and alpelisib led to a greater reduction in in AKT phosphorylation in the EGFR amplified cell lines (MDA-MB-468 and BT20) than either drug alone (Fig. 3B and D). MDA-MB-231 cells did not have a significant reduction in AKT signaling when comparing the drug combination to either drug alone (Fig. 3D and Fig. S3B). It should be noted that phosphorylated AKT levels were undetectable in MB231 at the exposure shown in Figure 3B, and the quantification of phosphorylated AKT levels in MB231 used in 3D were from immunoblots with longer exposures, such as the representative blot seen in Supplemental Figure S3B. All three cell lines had measurable p-P70S6K and pS6, however the drug combination only elicited a significant decrease in BT20 and MDA-MB-468 (Fig. 3B, E and F). In BT20 cells, afatinib or alpelisib alone reduced pS6 in a statistically significant manner, however the combination of the two drugs reduced pS6 to near undetectable levels (Fig. 3B and F). No significant effects on p-P70S6K or pS6 were seen in MDA MB-231. Similar results were observed using the EGFR TKI erlotinib and PI3K inhibitor BKM120 (Fig. S3B-F). Overall, this shows that the inhibition of pMAPK and a synergistic downregulation of pAKT, p- P70S6K and pS6K signaling occurs in response to dual EGFR and PI3K inhibition in cells with EGFR amplification and dysregulated PI3K. The greatest effect was observed in BT20 cells which have EGFR amplification and PIK3CA activating mutations.

**Figure 3.**
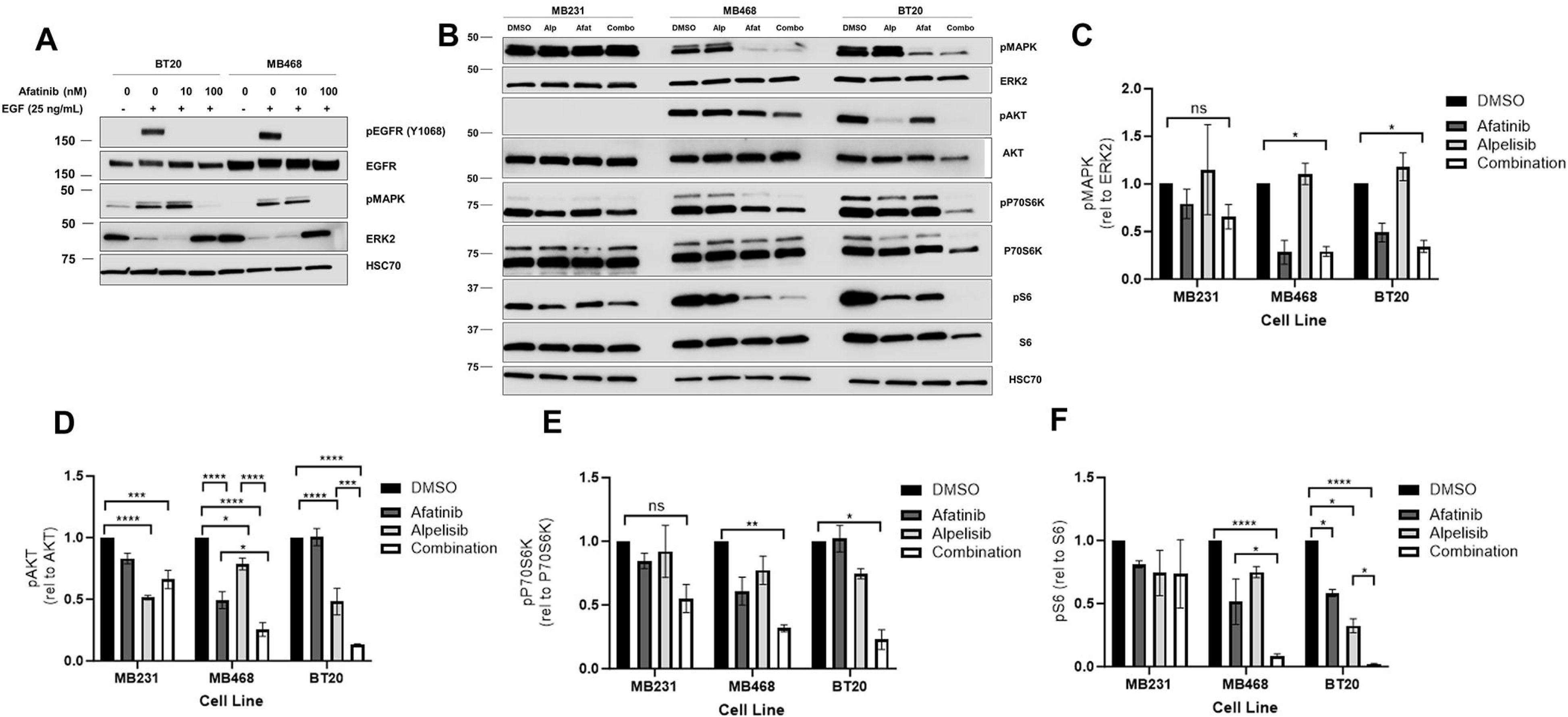
EGFR/PI3K dual inhibition significantly reduces downstream signaling in EGFR amplified and PI3K altered breast cancer. A) BT-20 and MDA-MB-468 cells were pretreated for 24 hours with 0, 10 or 100 nM afatinib, and then starved in serum-free RPM1 media for 3 hours prior to stimulation with or without 25 ng/mL EGF for fifteen minutes. Cells were lysed and immunoblotted for pEGFR (Y1068), EGFR, pMAPK (T202/Y204), ERK2 and HSC70 (loading control). B) MDA-MB-231, MDA-MB-468 and BT20 cells were treated with 100 nM alpelisib (Alp), 1 µM afatinib (Afat) or a combination of both for 24 hours, and were then lysed and immunoblotted for pAKT (S473), AKT, pMAPK (T202/Y204), ERK2, p-P70S6K (T389), P70S6K, pS6 (S235/236), S6 and HSC70 (loading control). In three independent experiments, the band density of the phosphorylated protein relative to total protein was averaged ±SEM for C) pMAPK/ERK2, D) pAKT/AKT, E) p-P70S6K/P70S6K and F) pS6/S6. Two-way ANOVA was performed, where ns indicates not statistically significant, * indicates p<0.05, ** indicates p<0.01, *** indicates p<0.001 and **** indicates p<0.0001.

It has been previously described that EGFR inhibition by cetuximab can increase expression of HER3, leading to heterodimerization of EGFR and HER3, in turn overriding the EGFR inhibition by cetuximab (55). We did not observe changes in HER3 or pHER3 levels after exposure to erlotinib. Further, the phosphorylation of HER3 was inhibited by erlotinib in the EGFR amplified cell lines (Fig. S4). HER3 levels were very low, and pHER3 phosphorylation was not detected in the MDA-MB-231 cells. It is likely that increased expression of HER3 cannot overcome inhibition by the TKI as it does for cetuximab since the kinase activity of EGFR is necessary for signaling of the EGFR/HER3 heterodimer (56).

### Dual EGFR and PI3K inhibition synergistically reduce cell viability in breast cancer cells with EGFR amplification and PI3K alteration

We next tested the effects of afatinib and alpelisib alone and in combination on cell viability. In MDA-MB-231 there was not significant viability inhibition by alpelisib alone, there was only an inhibition of viability at a high dose of afatinib, the combination did not significantly affect viability and synergy scores indicated an additive effect (Fig. 4A). In contrast, in BT20 and MDA-MB-468 cells, each drug alone significantly inhibited viability, but the combination showed statistical significance when comparing the combination of drugs to either drug alone, and exhibited synergy using all four calculations as determined by SynergyFinder (Fig. 4B and C) (48). Similar results were seen with erlotinib and BKM-120 (Fig. S5A-C). Overall, EGFR and PI3K inhibitors in combination exhibit a synergistic effect on cell viability in cell lines with EGFR amplification and PI3K alterations, and only an additive effect in a cell line without EGFR amplification or PI3K alterations.

**Figure 4.**
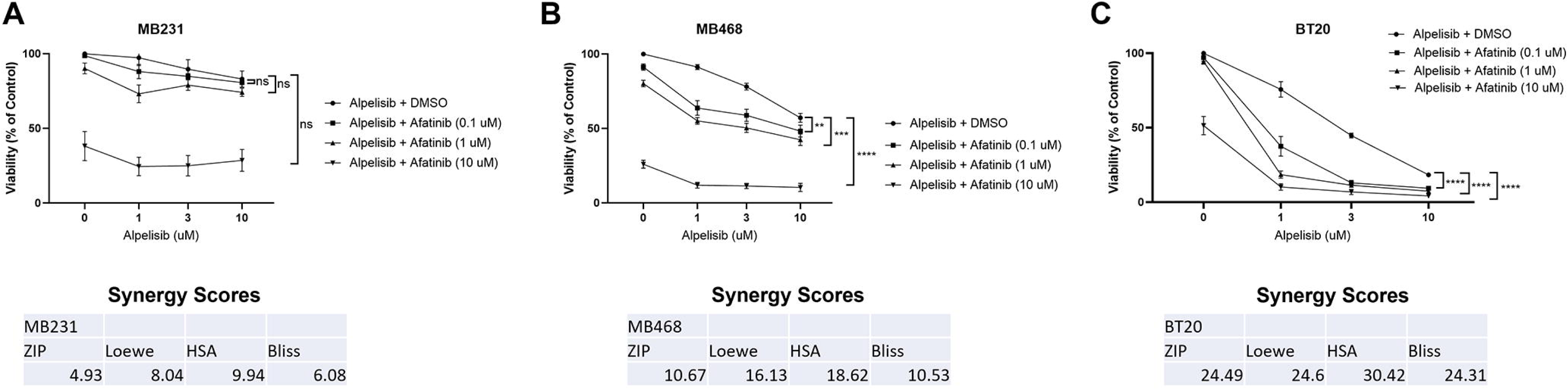
Dual EGFR/PI3K inhibition synergistically reduce cell viability in TNBC with EGFR amplification and PI3K alteration. A) MDA-MB-231 in three independent experiments, B) MDA-MB-468 in four independent experiments and C) BT20 cells in three independent experiments were treated with the indicated doses for 48 hours, and then viability was determined by Promega CellTiter-Glo 2.0. Data represents the average ±SEM of the independent experiments, and statistical analysis was performed as Two-Way ANOVA, and the p value indicates interaction, where ns indicates not statistically significant, ** indicates p<0.01, *** indicates p<0.001 and **** indicates p<0.0001. The averages were also subjected to four different synergy score analyses through synergyfinder.org, where a score of >+10 indicates synergy, -10 to +10 indicates additive effects, and <-10 indicates antagonism.

### Dual EGFR and PI3K inhibition induce cell cycle arrest and apoptosis in EGFR amplified and PI3K altered breast cancer cells

Cell cycle was not significantly altered in MB231 cells (Fig. 5A). In both MB468 and BT20, cell cycle analysis showed an increase in cells in the G1 phase of the cell cycle and statistically significant reductions in both the S phase and G2/M cell populations when EGFR and PI3K are inhibited by the combination of alpelisib and afatinib (Fig. 5B and C). Similar effects on the cell cycle were seen with erlotinib and BKM120, however there was also a statistically significant increase in sub-G1 population with the drug combination in both MDA-MB-468 and BT20 but not in MDA-MB-231 (Fig. S5 D-F).

**Figure 5.**
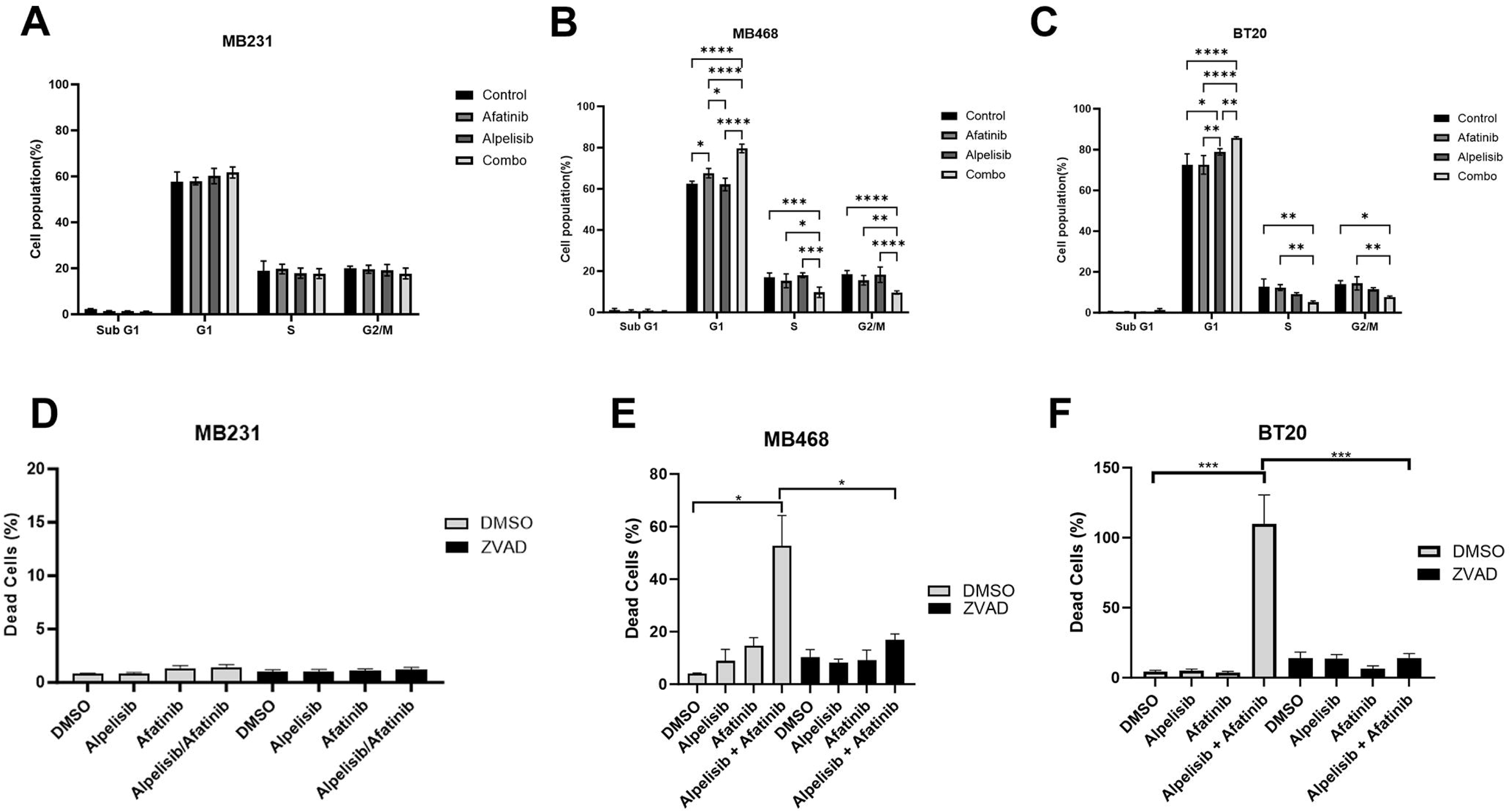
EGFR/PI3K inhibition induce cell cycle arrest and apoptosis in EGFR amplified and PI3K altered TNBC. A) MDA-MB-231 were treated with DMSO (control), 1 µM afatinib, 10 µM alpelisib, or the combination of both B) MDA-MB-468 were treated with DMSO (control), 1 µM afatinib, 10 µM alpelisib, or the combination of both C) BT20 were treated with DMSO (control), 100 nM afatinib, 1 µM alpelisib, or the combination of both and for 48 hours, and then were harvested, washed, fixed and stained with propidium iodide before being analyzed by flow cytometry and FlowJo for cell cycle changes. Data represents the average ±SEM of at least three independent experiments, and Two-Way ANOVA statistical analysis was performed where * indicates p<0.05, ** indicates p<0.01, *** indicates p<0.001 and **** indicates p<0.0001. D) MDA-MB-231 (1 µM alpelisib, 1 µM afatinib), E) MDA-MB-468 (3 µM alpelisib, 10 µM afatinib) or F) BT20 cells (1 µM alpelisib, 1 µM afatinib) were pretreated with 20 µM Z-VAD-FMK (ZVAD) for one hour, and then were treated with indicated drugs with continued ZVAD treatment in 1% FBS RPM1 media containing propidium iodide for 72 hours. Imaging of the cells was performed by BioTek Cytation 1, and dead cell percentage was calculated by dividing PI positive cells by the total brightfield cell count and multiplying by 100. Data represents the average ±SEM of three independent experiments. Statistical analysis was performed by Student’s t-test, with *** indicating p<0.001.

We quantified dead cell percentages using propidium iodide (PI) staining of cells treated with afatanib, alpelisib, and the combination in the presence or absence of the pan-caspase inhibitor Z-VAD-FMK (ZVAD). The combination of afatanib and alpelisib in both MB468 and BT20 cells resulted in a dramatic, statistically significant increase in dead cells which was rescued by ZVAD treatment, while the drugs alone or in combination did not increase the percentage of dead cells in MDA-MB-231 (Fig. 5D-F). Similarly, dual inhibition with erlotinib and BKM-120 increased the dead cells in BT20 and MDA-MB-468, which was rescued by ZVAD treatment (Fig. S6). Erlotinib and BKM-120 alone did increase caspase-dependent cell death in MDA-MB-468 and BT20 cells, but to a lesser extent (Fig. S6). MDA-MB-231 cells only showed a slight increase in cell death with erlotinib alone and no further increase in the percentage dead cells by the combination (Fig. S6A).

### Dual EGFR and PI3K inhibition reduce tumor volume *in vivo*

We conducted an orthotopic mammary fat pad xenograft model of BT20 cells in nude mice treated with control, afatanib, alpelisib or the combination (Fig 6A). Afatinib induced a statistically significant reduction in pEGFR when compared to controls and alpelisib induced a statistically significant reduction in pS6 compared to controls (Fig. 6B). Afatanib non- significantly reduced pS6 and alpelisib non-significantly reduced pEGFR. The combination of afatinib and alpelisib did not further reduce downstream signaling as measured by pS6 compared to alpelisib alone.

**Figure 6.**
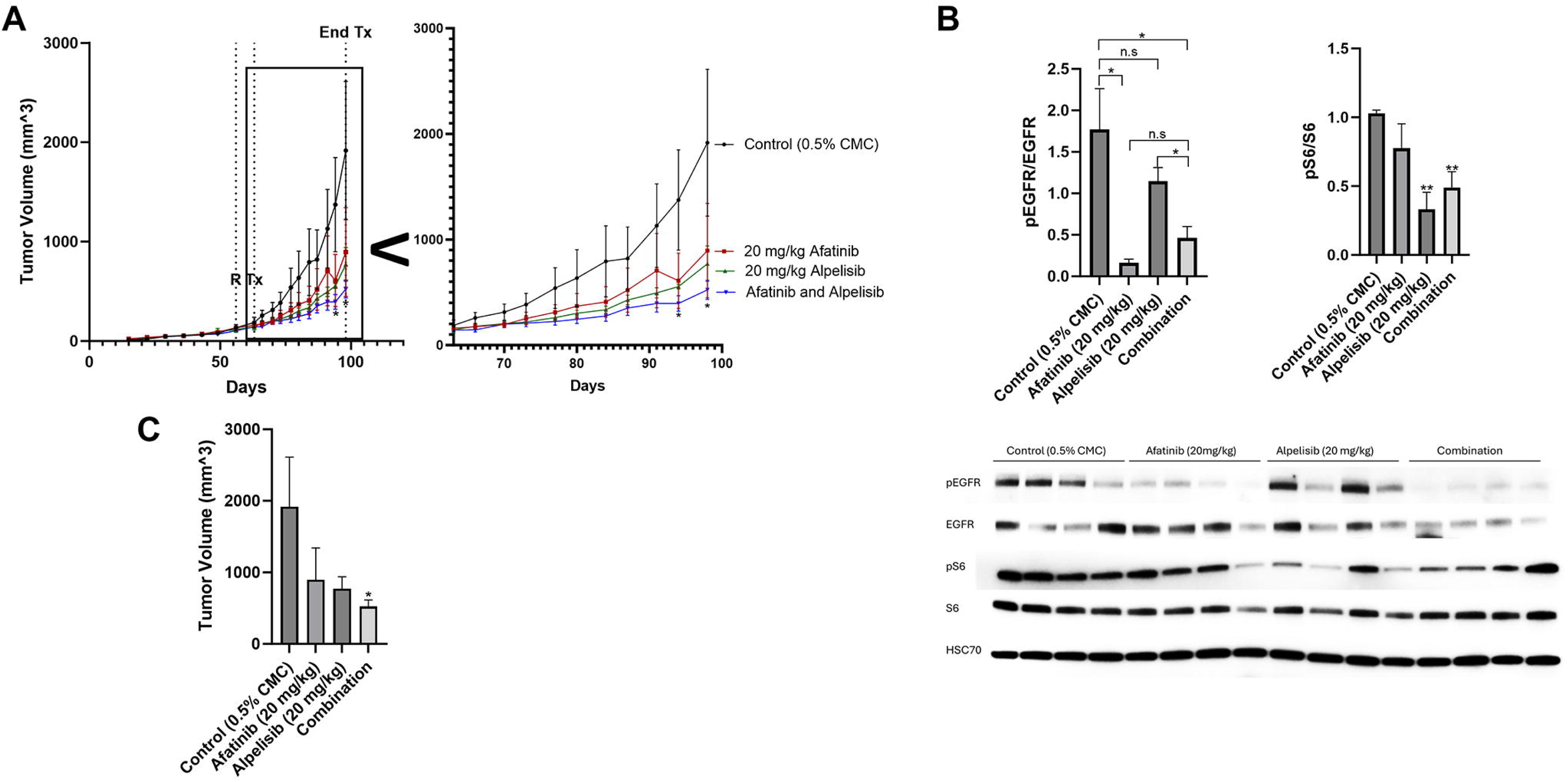
Dual EGFR and PI3K inhibition is most effective at reducing tumor volume *in vivo.* A) BT20 cells were injected subcutaneously into the mammary fat pad of nude mice. 56 days post implantation, when the average size was 100 mm^3^, mice were randomized (R) to 15 mice into four groups and treated starting on day 63 post implantation (Tx) with control (0.5% carboxymethylcellulose) (black), 20 mg/kg afatinib (red), 20 mg/kg alpelisib (green), or the combination of both afatinib and alpelisib (blue) Monday through Friday for 36 days. Zoomed inset shows day 63 to 100. Statistical analysis was performed by Student’s t-test, and * indicates p<0.05 when comparing afatinib and alpelisib to control on day 94 and day 98. B) After four days of treatment, four mice from each group were euthanized and the protein from the tumors was collected and immunoblotted for pEGFR (Y1068), EGFR, pS6 (S235/236), S6, and HSC70 (loading control). Data represents the average of four mice from each group ±SEM and statistical analysis was performed by Student’s t-test, where ns indicates not statistically significant, * indicates p<0.05 and ** indicates p<0.01. C) Bar graph of data from A on day 98 (36 days of drug treatment). Statistical analysis was performed by Student’s t-test, where * indicates p<0.05 when compared to control. No other comparisons were statistically significant.

Single agents alone reduced tumor volume after 36 days of treatment, however neither were statistically significant when compared to control due the slow and large variability in growth in vivo (Fig. 6A and C). The reduction in tumor size by the combination of both afatinib and alpelisib was statistically significant when compared to the control.

## Discussion

This studies goal was to determine the incidence of EGFR amplification in breast cancer patients and to examine the benefit of molecularly targeted agents in models of EGFR amplification. EGFR amplification occurs in approximately 1-5% of breast cancer patients with worse outcomes compared to patients without EGFR amplification (Table 1 and Fig. 1). This EGFR amplification was enriched in TNBC, HER2 amplified, and ER negative patients.

Importantly, TNBC patients are difficult to treat due to lack of defined molecular targets. Determination of targeted therapies aimed at a subset of breast cancer patients may provide therapeutic benefit in those who previously may have had poor prognosis and could help shift the paradigm to treating identifiable populations with a focused and specific molecular inhibition strategy.

Dual EGFR and PI3K inhibition in TNBC with EGFR amplification and PI3K aberrations act synergistically in reducing cell viability, as well as inducing apoptosis in vitro, and reducing tumor growth in vivo. Our data identifies a patient population which can be determined by molecular profiling and would benefit from treatment with this therapy. This study presents evidence for potential efficacy of EGFR and PI3K inhibitors in these patients and warrants clinical trial investigation into whether these patients would benefit from this therapy.

TNBC patients with EGFR amplification have a borderline worse survival compared to TNBC without EGFR amplification (Table 1 and Fig. S1A). Our data suggest that these patients may benefit from targeted EGFR and PI3K pathway inhibition. Further, this study characterized a model looking at TNBC, however ER+HER2- patients had a statistically significantly worse OS, therefore there are likely other breast cancer patient populations with EGFR amplification and PI3K alteration who might benefit from this dual therapy (Fig. S1).

The two cell lines with EGFR amplification and PI3K alterations (BT20 and MDA-MB-468) exhibited significant signaling pathway and biological effects of EGFR inhibitors and PI3K inhibitors while the MDA-MB-231 cells without the alterations were less affected by the drug combination (Fig. 3 and 4, Fig. S3). In a xenograft mouse model, the combination of EGFR and PI3K inhibitors reduced downstream signaling in the EGFR and PI3K pathways and inhibited the tumor growth in a statistically significant manner when compared to vehicle control (Fig. 6).

Dual EGFR and PI3K inhibition in vitro exhibited significant synergy in this study, however the in vivo data was more modest. The modest effects are likely due to the short half-life of alpelisib (approximately 1.5 hours in mice), whereas afatinib has a longer half-life (approximately 21 hours in mice) (57, 58). In the in vitro work cells were treated continuously for 48-72 hours with the drugs which could account for the difference in observed results when comparing in vitro and in vivo results in this study. The half-life of afatinib in human patients is approximately 30-40 hours and alpelisib is approximately 7.5 hours, suggesting with the daily dosing used there would be better steady state levels than in the mouse (59, 60).

Between 20-50% of breast cancers exhibit PIK3CA mutations, with the most common being hormone receptor positive or HER2 amplified breast cancer (61–66). It is noteworthy that PIK3CA alterations found in this study (∼36%) in EGFR amplified patients is similar to the percentage observed in HER2 amplified patients (30%) (64–66). Combining HER2 targeted therapy with PI3K inhibitors has been previously studied, and our study presents evidence that there is a similar population of EGFR amplified patients who could benefit from combination targeted therapy (67, 68).

EGFR and PI3K inhibitor combination has been studied in head and neck squamous cell carcinoma, glioblastoma, ovarian cancer and breast cancer preclinical cell line models, however these studies did not compare the synergistic effects of EGFR amplified and PI3K mutant cells to non-amplified and non-mutant cells (26–34, 36, 37). Further, a recent high throughput screen using breast cancer PDX models with PI3K mutations found that afatinib and alpelisib were synergistic, which confirms our data, however they did not compare the synergy of models with both EGFR amplification and PI3K mutations to those without (35). Our study indicates that it is important to determine the subset of patients who would benefit from this combination, by stratifying patients with EGFR amplification and PI3K pathway alterations. We also do not rule out the possibility of combining these drugs with HER2 targeted therapies, as we found that up to 7.7% of HER2 amplified/HER2 enriched tumors have amplified EGFR (Table 1). A study showing synergy with BKM120 and the HER2 inhibitor Trastuzumab in patients provides a proof of principle for combining PI3K inhibitors with EGFR family inhibitors may provide clinical benefit in patients with EGFR family amplification and PI3K mutation (69).

We studied multiple EGFR (erlotinib or afatinib) and PI3K (BKM120 or alpelisib) inhibitors (Fig. 3-6; Fig. S3, S5 and S6). Afatinib inhibits wild-type (WT) EGFR with the lowest IC50, and then at increasing IC50s it inhibits L858R EGFR, HER2 and HER4 (70, 71). In addition to PIK3CA, BKM120 also inhibits mTOR, VPS34 and DNAPK (72). Alpelisib is a more specific PIK3CA inhibitor, without observed off target effects. Since we tested different EGFR and PI3K inhibitors and observed similar reductions in downstream signaling, cell viability, cell cycle, and apoptosis when the drugs were combined, it is likely that the observed effects were due to a class effect of the drugs targeting EGFR and PIK3CA.

This study identifies that there is a subset of breast cancer patients with EGFR amplification with co-incident PI3K pathway mutations who are more likely to respond to this combination therapy. While this population represents a rare subset of metastatic breast cancer patients, they are identified by genomic analyses that are routinely performed on breast cancer patients as part of their clinical assessment. These analyses highlight approved drugs and potential clinical trials available to these patients but currently there are no clinical trial data or ongoing trials to support using EGFR and PI3K inhibitors in these patients and they are not treated with these agents. Therefore, the combination of EGFR and PI3K pathway inhibitors should be tested in breast cancer patients with EGFR amplification and PI3K pathway alteration. In order to test if there is activity, a phase I/II trial of 30-50 patients would likely be needed to see the response rate as this was sufficient to see activity of lapatinib in patients with metastatic HER2-amplified breast cancers (73). Currently, there are ∼150,000 patients living with metastatic breast cancer in the US (74). Based on our analysis shown in Table 1 and 2, with ∼1.2-5.1% of all breast cancers having an EGFR amplification, there would be ∼ 1800-7650 patients whose tumor has EGFR amplification, with approximately half having a mutation in the PI3K pathway. Gating only on TNBC, the prevalence of patients with metastatic TNBC is approximately 15% of the total, or about 22,500 cases (75). Based on our data showing the incidence of EGFR amplification between 3.2-8.4% in TNBC this would mean that that there would be approximately 720-1890 cases with EGFR amplification, again with about half having a PI3K pathway mutation. Thus, such a trial should be feasible. This ultimately could shift the paradigm for the treatment regimen in this patient population with EGFR amplification, especially triple negative breast cancer patients who have fewer targeted therapy options.

## List of Abbreviations

EGFR: Epidermal Growth Factor Receptor
PI3K: Phosphoinositide 3-kinase
TNBC: Triple Negative Breast Cancer
FISH: fluorescent in situ hybridization
PI: propidium iodide
ER: Estrogen Receptor
HER2: Human epidermal growth factor receptor 2
mTOR: mechanistic target of rapamycin
AKT: Protein Kinase B
EGF: epidermal growth factor
MAPK: mitogen activated protein kinases
TKI: tyrosine kinase inhibitor
P70S6K: ribosomal protein S6 kinase beta 1
SEM: standard error of the mean
TCGA: The Cancer Genome Atlas Program
MSKCC: Memorial Sloan Kettering Cancer Center
IDC: invasive ductal carcinoma
ILC: Invasive lobular carcinoma
OS: overall survival
PTEN: phosphatase and tensin homolog
DMSO: dimethyl sulfoxide
PDX: patient derived xenograft
WT: wild-type
DNAPK: DNA-dependent protein kinase

## Acknowledgements

We thank all members of the Lipkowitz lab in the Women’s Malignancies Branch, Center for Cancer Research, National Cancer Institute for their support and review of this manuscript. We would like to thank Dr. Thomas Ried for his assistance with the planning of the FISH experiment. We would like to acknowledge the CCR Genomics Core for their assistance with sequencing, and the veterinarians and animal care staff at NIH for their assistance with animal care, drug delivery and tumor measurement.

## Authors’ contributions

DJW performed immunoblotting, cell viability assays, cytotoxicity assays, xenograft assays, helped with project conceptualization, experimental design, data analysis and interpretation, and wrote the manuscript. DV performed immunoblotting, cell viability assays, cytotoxicity assays, prepared samples for FISH and sequencing, performed flow cytometry, helped with project conceptualization and experimental design. YA performed immunoblotting and flow cytometry, helped with experimental design and data analysis. SD analyzed Caris datasets and helped with data analysis and interpretation. SW ML and GS assisted with the Caris dataset. Darawalee Wangsa, Danny Wangsa and KH performed FISH analysis. YEG helped with experimental design, data analysis and interpretation, and animal studies. SL analyzed cBioPortal datasets, helped with project conceptualization, data analysis and interpretation, and manuscript editing. All authors read and approved the final manuscript.

## Ethics approval and consent to participate

All animal aspects of this study were approved by the NCI Animal Care and Use Committee (IACUC number WMB-004). All studies in the Caris dataset were conducted in accordance with guidelines of the Declaration of Helsinki, Belmont report, and U.S. Common rule, and in keeping with 45 CFR 46.101(b)(4), was performed utilizing retrospective, deidentified clinical data. Per WCG IRB, it was considered institutional review board (IRB) exempt with waiver of patient consent. 15,247 breast cancer samples underwent molecular testing at Caris Life Sciences (Phoenix, AZ, USA), a CLIA/CAP-certified laboratory. No specific inclusion criteria or exclusion criteria were applied aside from the requirement for successful sequencing results.

## Data Availability Statement

The MSKCC, TCGA and METABRIC datasets analyzed in this study are available on cBioPortal.org (41–43). The Caris deidentified sequencing data are owned by Caris Life Sciences and cannot be publicly shared due to the data usage agreement in place. These data will be made available to researchers for replication and verification purposes through the Caris letter of intent process, which are generally fulfilled within 6 months. For more information on how to access this data, please contact Joanne Xiu at. All other data generated in this study are included in this published article and its supplementary information.

## Supplemental Figures

**Figure S1**

File Format: .tiff

Title of Data: Overall survival of patients with EGFR amplification by subtype.

Figure S1 Legend: Using the Caris dataset, overall survival of breast cancer patients with EGFR amplification (red) vs WT (blue) in A) Triple negative breast cancer, B) Basal, C) HER2 enriched, or D) ER+ HER2-.

**Figure S2**

File Format: .tiff

Title of Data: Oncoplot for PI3K pathway alterations in EGFR amplified breast cancer.

Figure S2 Legend: Patient data from cbioportal.org plotting varying mutations or amplifications by gene in the PI3K pathway.

**Figure S3**

File Format: .tiff

Title of Data: EGFR/PI3K dual inhibition significantly reduce downstream signaling in EGFR amplified and PI3K altered TNBC.

Figure S3 Legend: A) MDA-MB-468 and BT20 cells were starved in serum-free RPM1 media for 1 hour with 0, 1 or 10 µM erlotinib, and then stimulated with or without 25 ng/mL EGF for 10 minutes. Cells were lysed and immunoblotted for pEGFR (Y1068) and EGFR (loading control). B) Darker exposure for MDA-MB-231 panel from Figure 4B which was used for quantification. C) MDA-MB-231, MDA-MB-468 and BT20 cells were treated with 10 µM erlotinib, 1 µM BKM120 or a combination of both for 24 hours, and were then lysed and immunoblotted for pAKT (S473), AKT, pMAPK (T202/Y204), ERK2, p-P70S6K (T389), P70S6K and HSC70 (loading control). In three independent experiments, the band density of the phosphorylated protein relative to total protein was averaged ±SEM for D) pMAPK/ERK2, E) pAKT/AKT and F) p-P70S6K/P70S6K. Student’s t-test was performed, where ns indicates not statistically significant, * indicates p<0.05, ** indicates p<0.01, *** indicates p<0.001 and **** indicates p<0.0001.

**Figure S4**

File Format: .tiff

Title of Data: Erlotinib treatment does not activate pHER3.

Figure S4 Legend: MDA-MB-231, MDA-MB-468 and BT20 cells were treated with 10 µM erlotinib, 1 µM BKM120 or the combination of both for 24 hours before the cells were lysed and immunoblotted for pHER3 (Y1289) and HER3 (loading control).

**Figure S5**

File Format: .tiff

Title of Data: EGFR/PI3K inhibition reduces viability, induces cell cycle arrest in TNBC with EGFR amplification, PI3K alteration.

Figure S5 Legend: A) MDA-MB-231, B) MDA-MB-468 and C) BT20 cells were treated with the indicated doses for 48 hours, and then viability was determined by Promega CellTiter-Glo 2.0. Data represents the average ±SEM of three independent experiments, and statistical analysis was performed as Two-Way ANOVA, and the p value indicates interaction, where ns indicates not statistically significant, * indicates p<0.05, *** indicates p<0.001 and **** indicates p<0.0001. D) MDA-MB-231, E) MDA-MB-468 and F) BT20 cells were treated with DMSO control, 1 µM erlotinib or 10 µM BKM120 or the combination of erlotinib and BKM120 for 48 hours, and then were harvested, washed, fixed and stained with propidium iodide before being analyzed by flow cytometry and FlowJo for cell cycle changes. Data represents the average ±SEM of at least three independent experiments, and Two-Way ANOVA statistical analysis was performed where * indicates p<0.05, ** indicates p<0.01 and *** indicates p<0.001.

**Figure S6**

File Format: .tiff

Title of Data: Dual EGFR/PI3K inhibition induce apoptosis in EGFR amplified and PI3K altered TNBC.

Figure S6 Legend: A) MDA-MB-231, B) MDA-MB-468 and C) BT20 cells were pretreated with 20 µM Z-VAD-FMK (ZVAD) for one hour, and then were treated with DMSO, 10 µM erlotinib, 1 µM BKM120 or the combination of both, with continued ZVAD treatment for 48 hours. Dead cell percentage was calculated by using CytoTox-Glo by Promega. Data represents the average ±SEM of three independent experiments. Statistical analysis was performed by One-way ANOVA, with ns indicating not statistically significant, * indicating p<0.05, ** indicating p<0.01 and *** indicating p<0.001.

