## Supplementary figures and images for "EGFR amplification and PI3K pathway mutations identify a subset of breast cancers that synergistically respond to EGFR and PI3K inhibition"

### Supplemental Figure S1

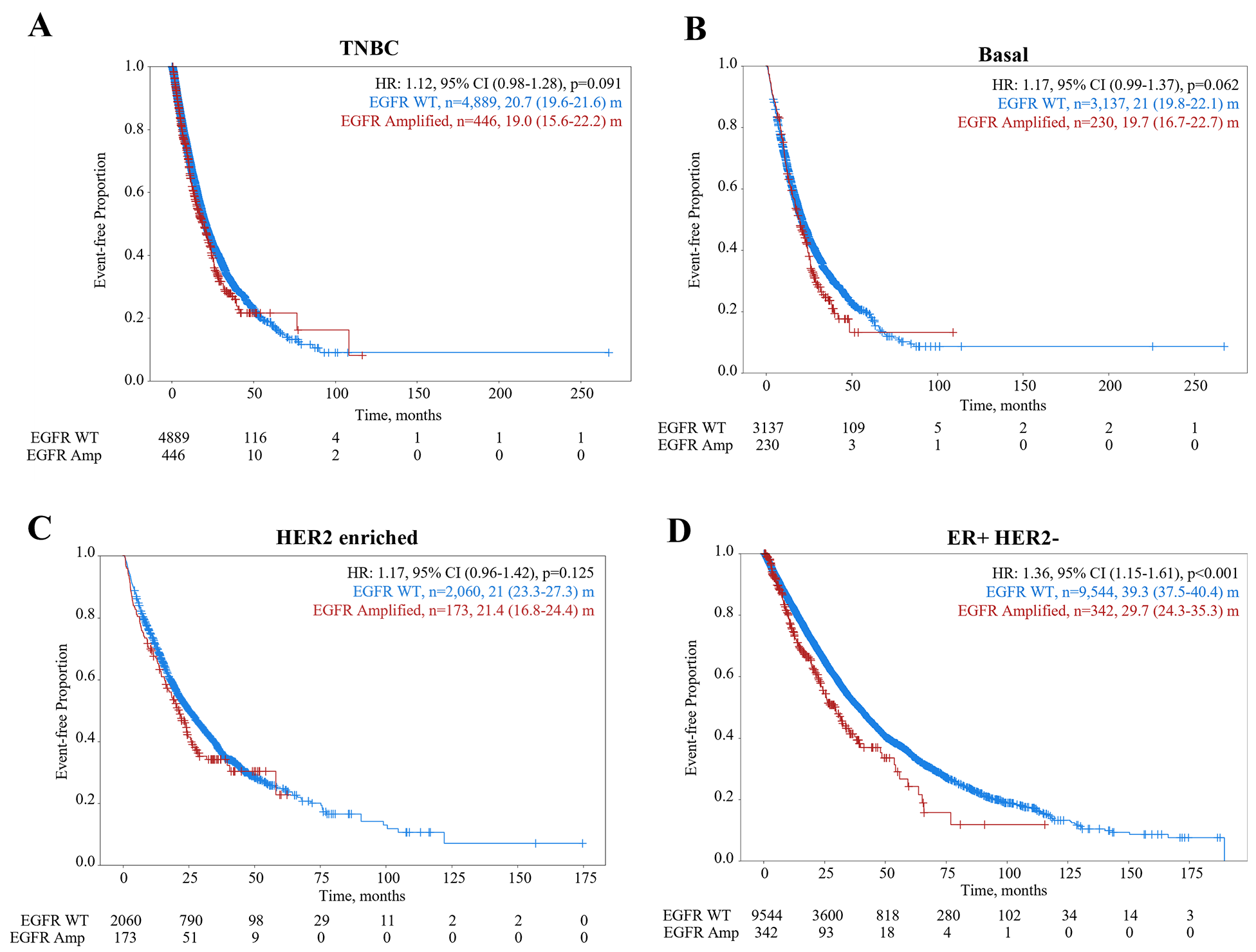

### Supplemental Figure S2

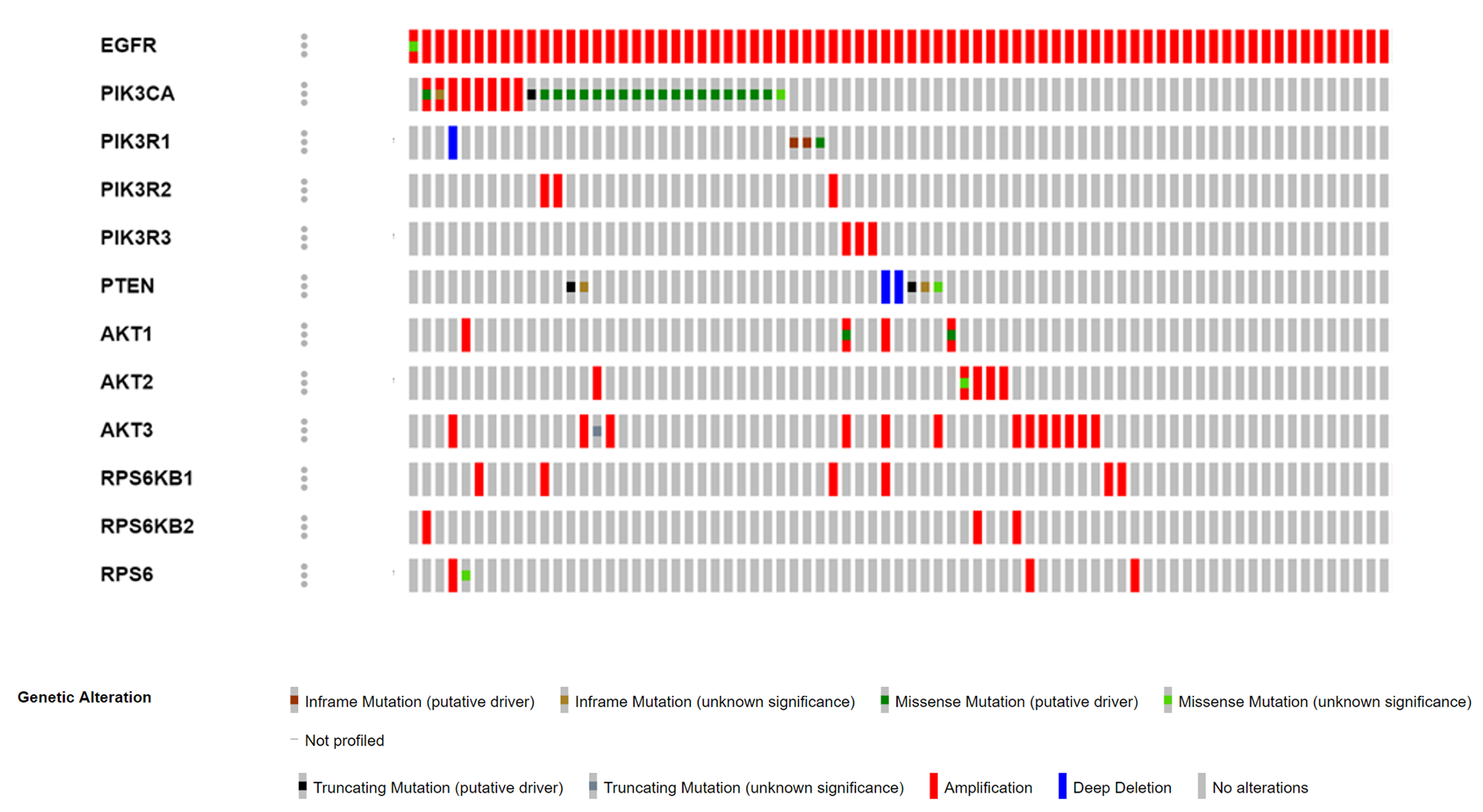

### Supplemental Figure S3

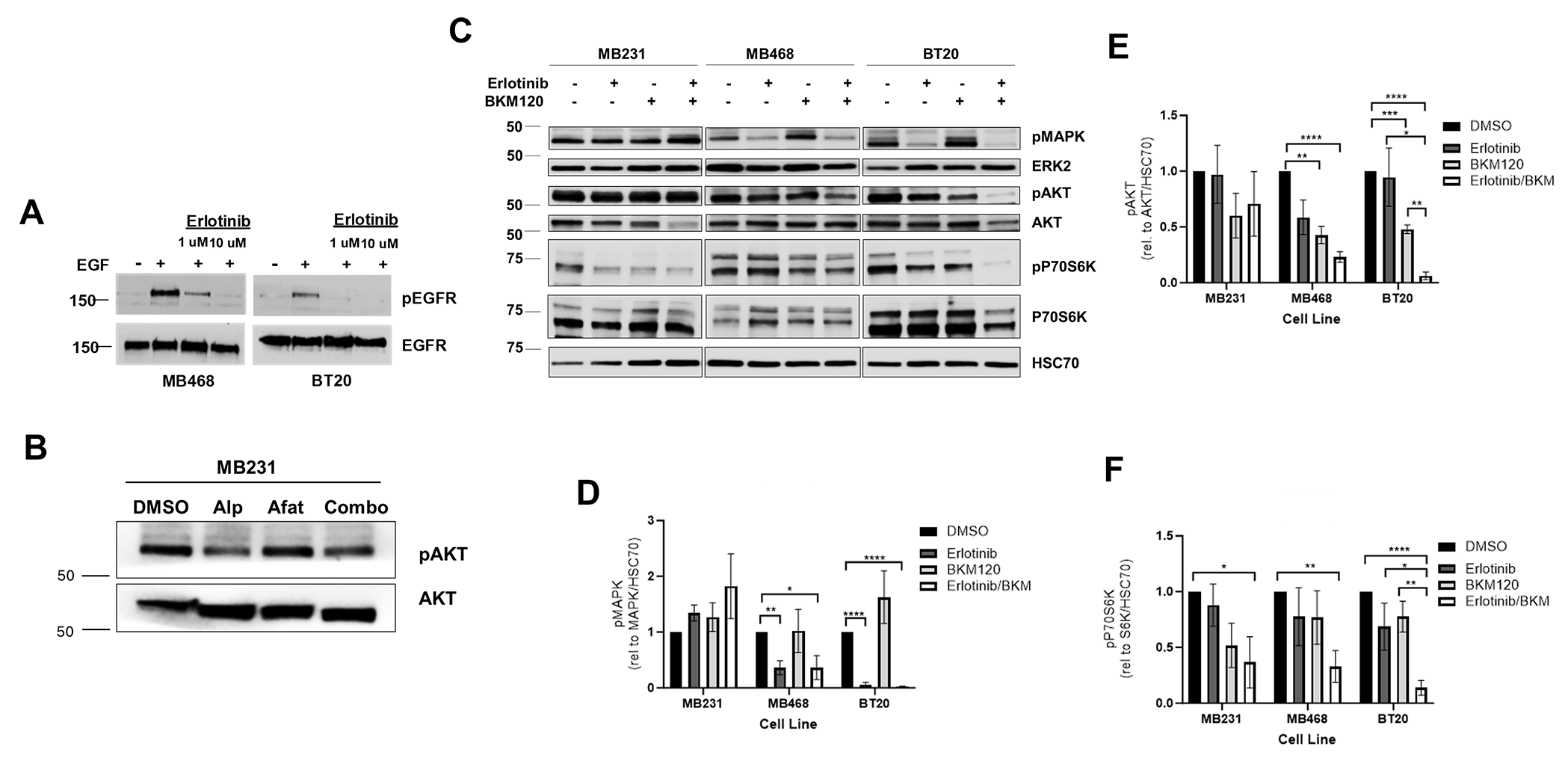

### Supplemental Figure S4

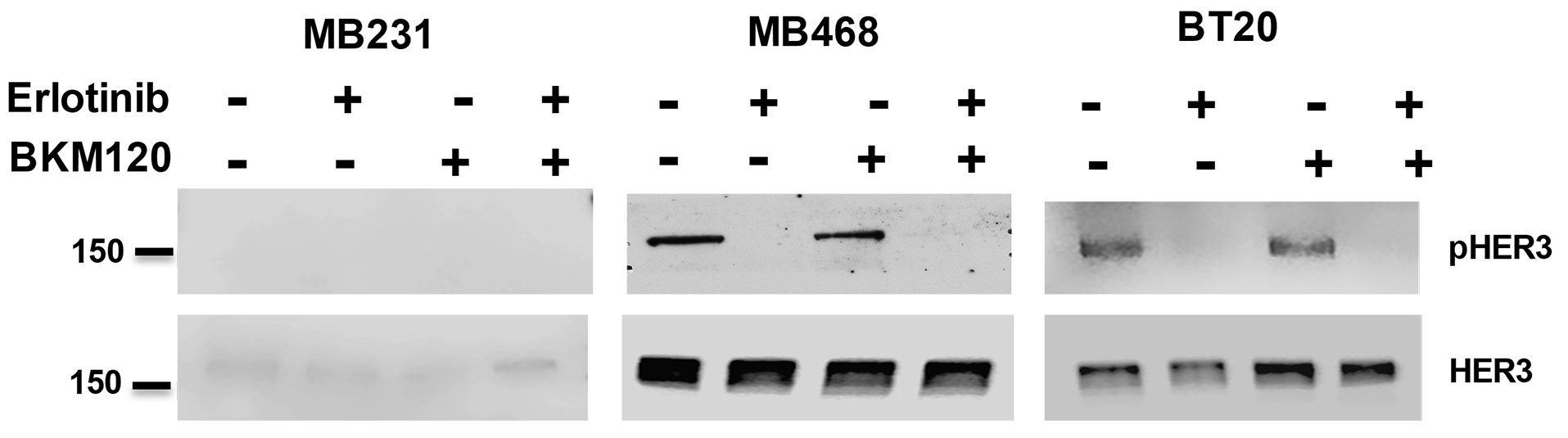

### Supplemental Figure S5

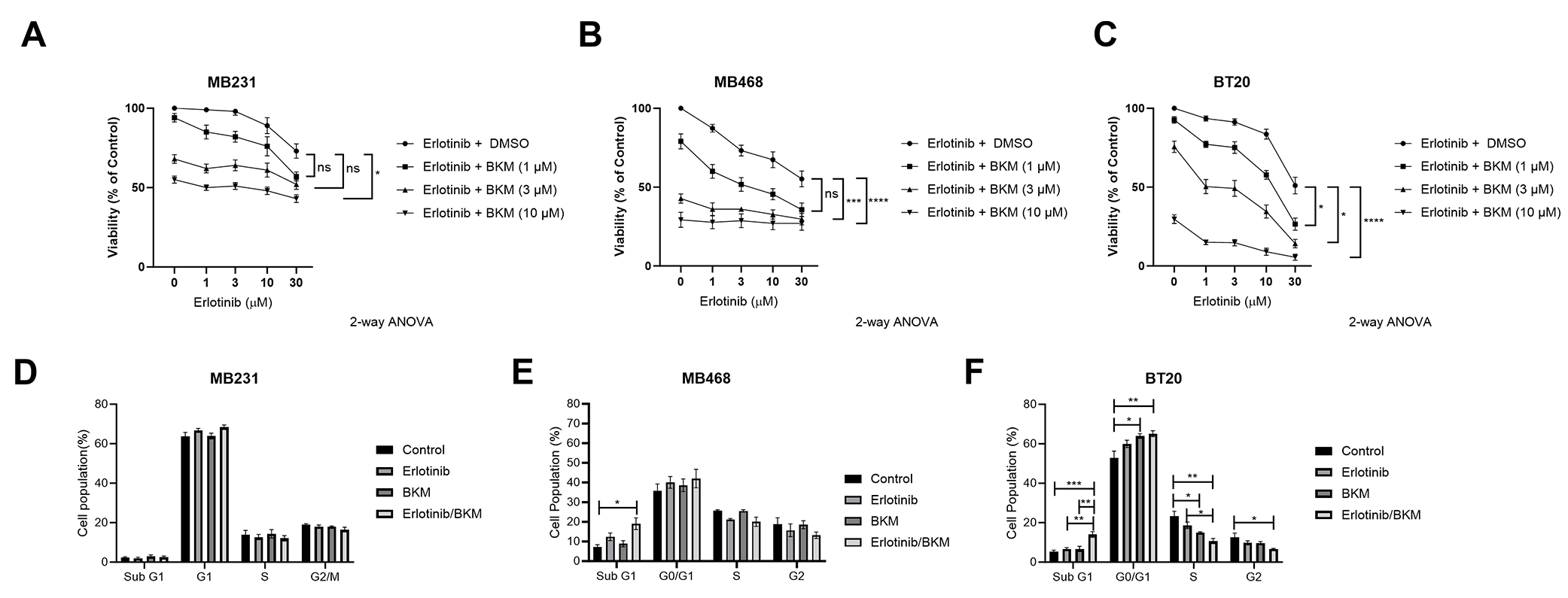

### Supplemental Figure S6

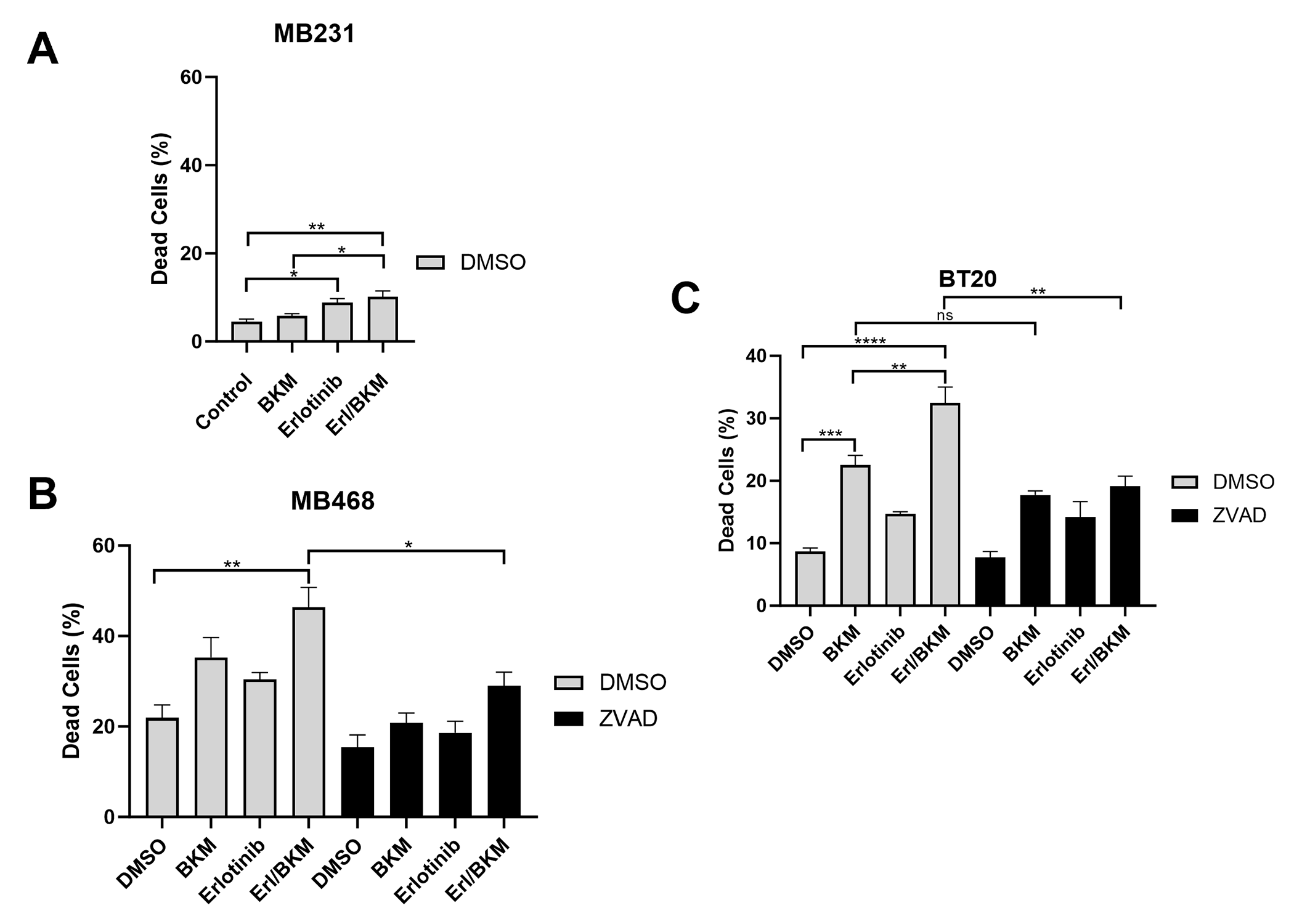
